# Glycolytic ATP production enables rapid cell growth

**DOI:** 10.1101/2025.10.30.685682

**Authors:** Matthew A. Kukurugya, Shuying Zhang, Brandon T. Ha, Alex E. Ekvik, Denis V. Titov

## Abstract

Cells use ATP to fuel growth and maintenance. Surprisingly, much of the ATP in rapidly growing cells is produced through glycolytic fermentation rather than the more efficient process of respiration, a puzzling phenomenon known as the Warburg Effect. One proposed explanation is that glycolysis produces ATP faster than respiration, thereby enabling faster growth rates. But whether this explains the Warburg Effect is uncertain, as the ATP costs of mammalian cell growth have not been rigorously estimated. Here, we perform over 7500 measurements to estimate the ATP costs of growth and maintenance in mammalian cells across a range of growth rates and under perturbations to ATP production and demand. We find that respiration alone cannot meet the ATP demands of cells doubling faster than every 30 hours, roughly one-third of the maximal mammalian cell growth rate, demonstrating that the Warburg Effect is required to sustain the ATP demands of rapid cell growth.

## Introduction

Rapidly proliferating cells produce a large fraction of ATP through glycolytic fermentation rather than the more efficient oxidative phosphorylation, even in the presence of abundant oxygen, a phenomenon known as the Warburg Effect^1,2^. The Warburg Effect is puzzling, as it is unclear why cells would prefer to use glycolysis in the presence of oxygen, even though it produces 10 times less ATP per glucose than respiration. Yet, this preference for glycolysis is one of the most reproducible metabolic features of proliferating cells, observed across cancer cells^1,2^, activated lymphocytes^3,4^, embryonic stem cells^5,6^, *S. cerevisiae*^7–9^ and even *E. coli*^10–13^. Many hypotheses to explain the Warburg Effect have been proposed^14–20^, but none are universally accepted.

We and others have proposed that the Warburg Effect is the result of cells utilizing glycolysis to maximize their ATP production rate when ATP demand is high and glucose is abundant^9,11,21–25^. This hypothesis requires two conditions to be satisfied. First, glycolysis must produce ATP faster than respiration per mg of pathway protein, as has been shown for mammalian cells^25^, *S. cerevisiae*^9,24,25^, and *E. coli*^11,25^. In other words, if cells dedicate a fraction of their proteome to ATP-producing enzymes, they’ll produce ATP at a faster rate if that fraction is occupied by glycolytic enzymes. Second, ATP production must be limiting for rapid growth, such that respiration alone is not fast enough to support the ATP demands of rapidly growing cells. The aim of this study is to test whether the second condition holds by rigorously quantifying the ATP costs of mammalian cell growth.

The ATP requirements of cell growth have been extensively studied in microbes, where the ATP production rate increases linearly with growth rate^27,28,32^. The linear relationship enabled the decomposition into ATP required for cell growth and cell maintenance without growth. For *E. coli* and *S. cerevisiae* growing near maximal rates, the majority of ATP was used for growth rather than maintenance. On the contrary, the two studies in mouse LS cells concluded that the majority of ATP was used for cellular maintenance and that ATP production increased only modestly with growth rate. These two studies have been cited^15–17^ as evidence that ATP demand is not limiting for mammalian cell growth, but rely on only a few measurements from a single slow-growing cell line.

Here we systematically measure the ATP costs of growth and maintenance across twelve mammalian cell lines. Using over 7,500 measurements of glycolytic rates, respiratory rates, cell sizes, and growth rates, combined with four orthogonal perturbations of ATP supply and demand, we show that ATP production scales linearly with growth rate in mammalian cells, that growth-associated costs account for the majority of ATP expenditure in rapidly dividing cells, and that ATP production capacity causally limits growth rate. We further show that the glycolytic fraction of ATP production increases with growth rate across mammalian cells, *S. cerevisiae* and *E. coli*. Finally, we show that respiration alone is insufficient to sustain doubling times faster than approximately 30 hours in mammalian cells, roughly one-third of the maximal mammalian cell growth rate. These results overturn the prior consensus on mammalian energy allocation, establish a causal role for ATP supply in limiting proliferation, and provide a quantitative and mechanistic explanation for why rapidly growing mammalian cells require the Warburg Effect.

## Results

### ATP production scales linearly with growth rate in mammalian cells and growth dominates ATP expenditure

We measured ATP production rate and growth rate across twelve mammalian cell lines spanning a range of cell sizes, growth rates, and tissue origins (Fig. 1a, Fig. S1). ATP production rate was calculated as the sum of glycolytic and respiratory contributions, using lactate production and oxygen consumption rates multiplied by their respective ATP yields (Fig. S1, also see Method S1. Detailed calculation of cellular ATP production rates from glycolysis and respiration). Each value was averaged from three independent measurements, with several biological replicates for each, totaling around 200 measurements per point in Fig. 1a. This represents the largest single compilation of ATP production measurements for mammalian cells. We observed a strong linear relationship between ATP production rate and growth rate across all twelve cell lines in the presence of various perturbations described below (Fig. 1a; R^2^ = 0.72, p = 5.6 10⁻^11^), consistent with data for *E. coli* and *S. cerevisiae* (Fig. 1b,c). The slope of the linear fit estimates the ATP cost of producing one milligram of new cell protein (growth cost, *C*_growth_), and the y-intercept estimates the ATP production rate required to sustain non-dividing cells (maintenance cost, *V*_maint_), related by

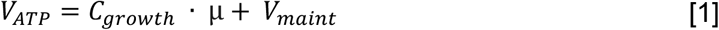

**Fig. 1.**
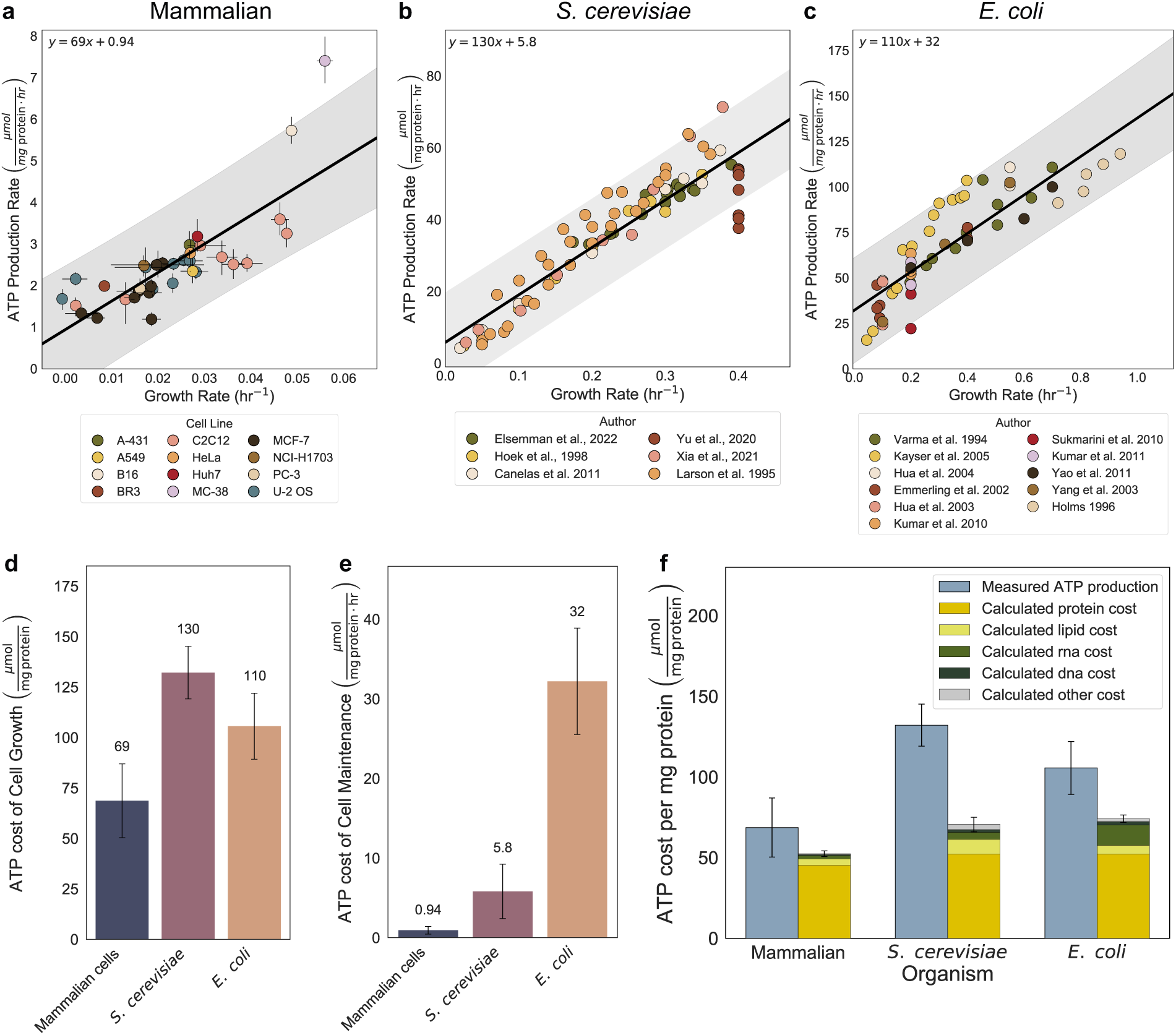
ATP production rate scales linearly with growth rate across mammalian cells, *S. cerevisiae*, and *E. coli*, and growth, not maintenance, dominates ATP expenditure. (a–c) ATP production rate (µmol mg protein⁻¹ hr⁻¹) versus growth rate (hr⁻¹) in (a) mammalian cells across twelve cell lines, (b) *S. cerevisiae*, and (c) *E. coli*. In (a), point colors denote cell line; in (b) and (c), point colors denote data source (study), compiled from published datasets. Each point is an individual measurement. The solid line is the ordinary least-squares linear fit (equation inset); the shaded band is the 95% prediction interval from 10,000 bootstrap iterations. Error bars represent 95% confidence intervals. (d) Slope of the linear ATP–growth fit (ATP cost of cell growth, µmol ATP per mg protein) per organism (e) Y-intercept of the fit (ATP cost of cell maintenance, µmol mg protein⁻¹ hr⁻¹) for per organism. (f) Measured ATP production (slope from a–c; blue bar) per organism compared side-by-side with the total theoretical biosynthetic ATP cost, the latter shown as a stacked bar broken down by macromolecular class (protein, lipid, RNA, DNA, other). Error bars in (d–f) represent 95% confidence intervals from bootstrap resampling.

Estimated growth costs were 69, 130, 110 µmol ATP per mg protein for mammalian cells, *S. cerevisiae*, and *E. coli*, respectively (Fig. 1d). These values are broadly conserved across organisms, reflecting the shared ATP costs of peptide elongation and nucleic acid synthesis; the lower mammalian value reflects nutrient-replete culture conditions in which amino acids and lipids are supplied exogenously. Estimated maintenance costs were 0.94, 5.8, and 32 µmol ATP per mg protein per hour for mammalian cells, *S. cerevisiae*, and *E. coli*, respectively (Fig. 1e), consistent with Kleiber’s law scaling of metabolic rate with cell size^36^. Critically, at a doubling time of 24 hours, growth-associated ATP consumption accounts for approximately 75% of total ATP production in mammalian cells, directly contradicting prior reports that maintenance dominates ATP expenditure^34,37,38^. Comparison of measured growth costs to theoretical biosynthetic costs suggests that protein synthesis is the dominant driver of ATP demand during growth and that biosynthesis accounts for 77 percent, 54 percent, and 70 percent of measured ATP production in mammalian cells, *S. cerevisiae*, and *E. coli* (Fig. 1f, Fig. S2).

We compared our estimates with literature values, when available, to further validate our methodology (see Method S5: Conversion of published values to common units for discussion of unit conversion for literature values). Our values for MCF-7 cells in Fig. 1a, reported as mean ± SD, are in good agreement with previously reported measurements 0.018 ± 0.005 vs. 0.015 ± 0.003 h⁻¹ for growth rate, 0.54 ± 0.22 vs. 0.77 ± 0.14 for lactate production, and 0.32 ± 0.06 vs. 0.64 ± 0.01 for oxygen consumption^39^. Notably, the previously reported oxygen consumption rates were not corrected for non-mitochondrial oxygen consumption^39^. For *E. coli* and *S. cerevisiae*, our measured growth costs of 110 and 130 µmol ATP per mg protein and maintenance costs of 32 and 5.8 µmol ATP per mg protein per hour, together with the calculated biosynthetic costs of 74 and 71 µmol ATP per mg protein, agree with both the measured and the calculated values reported previously^28,40–43^. For mammalian cells, the only prior estimates^34,35^ came from slowly growing mouse LS cells reported growth costs of 51 and 220 and maintenance costs of 1.4 and 7.9 µmol ATP per mg protein per hour, both assuming an ATP yield of 38 per glucose well above our value of 24, with the first study being close to our measured 69 and 0.94. Our estimates for maintenance cost are also similar to basal metabolic rate in humans where a metabolic equivalent of 3.5 ml O2 per kg wet weight per min^44^ which equates to 0.25 µmol ATP per mg protein per hour. After accounting for metabolically inert connective tissue protein (e.g., collagen is estimated to be up to 40% of protein in the human body^45,46^), the effective maintenance cost per mg of metabolically active protein in the human body is within 2-fold of our estimate for mammalian cells in culture.

### ATP production capacity causally limits mammalian cell growth

To test whether the ATP–growth correlation is causal, we performed four orthogonal perturbations of ATP supply and demand, asking in each case whether perturbed cells fall on the same linear relationship as unperturbed cells (Fig. 2). We inhibited glycolysis by targeting lactate dehydrogenase (LDH) with GSK-2837808A (GSK; IC₅₀ = 2.6 nM for LDHA, 43 nM for LDHB; 0–150 µM) in MCF-7, U-2 OS, and C2C12 cells. GSK treatment produced dose-dependent decreases in lactate production, total ATP production, and growth rate, with perturbed cells falling on the same linear relationship as unperturbed cells (Fig. 2b). Oxygen consumption increased modestly but was insufficient to compensate for the loss of glycolytic ATP, indicating that respiratory capacity was already near its limit in these cells (Fig. S3). We then inhibited Complex III of the electron transport chain with antimycin A, acutely suppressing oxidative phosphorylation. Even with respiratory ATP production abolished, the linear ATP–growth relationship was preserved, because cells fully compensated by upregulating glycolysis (Fig. 2d). Strikingly, growth rates remained high under antimycin treatment, demonstrating that glycolysis alone is sufficient to sustain rapid proliferation and that respiration is dispensable for growth when glycolytic capacity is intact (Fig. S3). Respiration could not compensate for glycolytic inhibition, but glycolysis fully compensated for respiratory inhibition, directly demonstrating that respiratory capacity close to saturation at baseline while glycolytic capacity retains sufficient reserve. We next inhibited protein synthesis with cycloheximide (0–1 µM), which blocks eukaryotic translation elongation and directly reduces ATP demand. Cycloheximide produced dose-dependent decreases in growth rate accompanied by proportional reductions in total ATP production, driven primarily by reduced lactate production (Fig. 2f, Fig. S3). Together, these three perturbations establish a bidirectional causal relationship in which ATP production rate constrains growth and growth rate dictates ATP production requirements, with glycolysis as the dominant supply pathway.

**Fig. 2.**
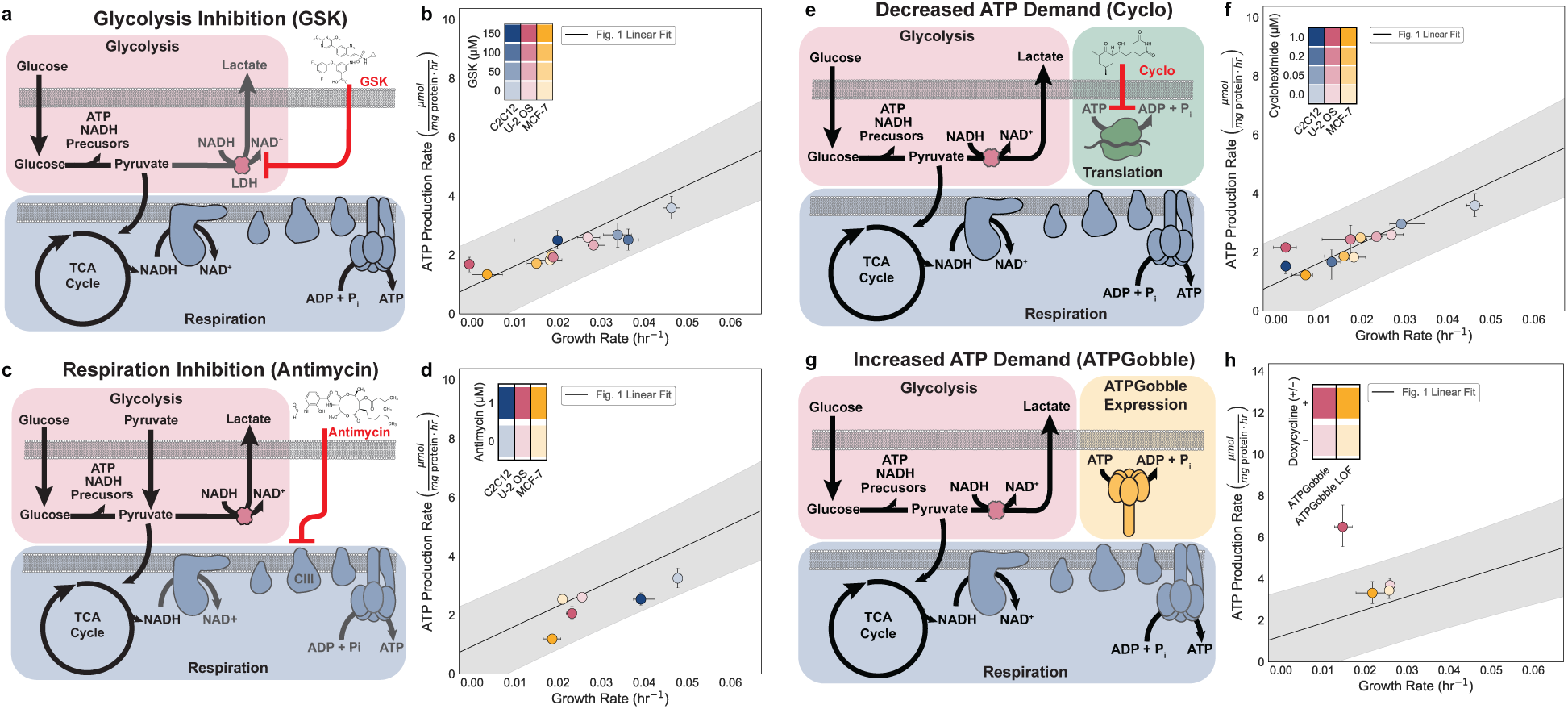
ATP production capacity causally limits mammalian cell growth. (a–d) Growth rate and total ATP production rate in MCF-7, U-2 OS, and C2C12 cells following inhibition of glycolysis with GSK-2837808A (0–150 µM), inhibition of respiration with antimycin A (b; 0 or 1 µM), inhibition of translation with cycloheximide (0–1 µM), or induction of ATPGobble in U-2 OS cells ( +DOX vs. loss-of-function control). In all panels, the solid line and shaded band show the linear fit and 95% prediction interval from Fig. 1a. Points represent group means; colors denote cell lines; shading indicates inhibitor dose or condition. Error bars show 95% CIs from 10,000 bootstrap resamples.

To test whether ATP production capacity sets an absolute ceiling on growth, we expressed ATPGobble, a genetically encoded ATP sink consisting of a modified *E. coli* F₁-ATPase that hydrolyzes ATP without performing useful work^47^. Unlike pharmacological inhibitors, ATPGobble increases ATP demand without directly perturbing any metabolic pathway, providing a direct test of whether cells can upregulate ATP production to compensate for an imposed increase in ATP demand. Induction of ATPGobble in U-2 OS cells increased total ATP production, with marked increases in both lactate production and oxygen consumption, yet growth slowed by approximately 2-fold relative to uninduced and loss-of-function controls (Fig. 2h, Fig. S3). The fact that cells upregulated both glycolysis and respiration in response to ATPGobble, yet still could not maintain their original growth rate, demonstrates that the cell was already operating near the ceiling of its total ATP production capacity. This result rules out the alternative interpretation that the ATP–growth correlation reflects demand-driven regulation alone and establishes that supply capacity is a limiting factor for mammalian cell proliferation.

### The glycolytic fraction of ATP production increases with growth rate across species

We next investigated how glycolysis and respiration contribute to total ATP production as a function of growth rate. In microbes, it is well established that cells switch from exclusively using respiration to a mix of glycolysis and respiration as growth rate increases^7,11^. Previous measurements in mouse LS cells^34,48^ and activated T cells^49^ suggest that mammalian cells exhibit the same phenomenon. In addition, many tumors upregulate glucose transporters and key glycolytic enzymes, supporting high glycolytic flux even with replete oxygen^14,50^. We observed that in our data for mammalian cells, the fraction of ATP produced by glycolysis increases with growth rate while that of respiration decreases (Fig. 3a), and a similar relationship was observed for microbes (Fig. 3b,c). Notably, *E. coli* and *S. cerevisiae* remain predominantly respiratory until respiratory ATP production reaches its growth-limiting capacity, after which glycolysis is required to support further increases in growth rate (Fig. 3b,c). A piecewise-linear fit to the data in Fig. 3b,c clearly delineates these two regimes in *E. coli* and *S. cerevisiae*: a respiratory regime in which respiration alone fuels slow growth, and a glycolytic regime in which glycolysis supplements respiration to fuel rapid growth. On the contrary, a piecewise-linear fit to the mammalian cell data in Fig. 3a produces only one regime of mixed glycolysis and respiration, suggesting that proliferating mammalian cells are typically in the state primed for maximal ATP production and growth rates. Across all three organisms, glycolysis accounts for about half of total ATP production at higher growth rates (Fig. 3), supporting the hypothesis that glycolysis plays a critical role in meeting the elevated ATP demands of rapidly proliferating cells. Review of the absolute values of glycolysis and respiration rate shows that the change in relative contribution of each pathway is almost exclusively due to a dramatic increase in glycolysis rate (Fig. 3d-f), while respiration rate is approximately flat (Fig. 3g-i). A piecewise-linear fit of the absolute rates for each pathway again clearly shows two regimes for microbes and only one for mammalian cells (Fig. 3d-i). A previous study found that the glycolytic rate did not correlate with growth rate in mammalian cells^51^; however, this interpretation was influenced by the use of glycolytic rate per cell, whereas using biomass as the normalization metric shows a clear correlation between glycolytic activity and growth, as has been pointed out by others^52^.

**Fig. 3.**
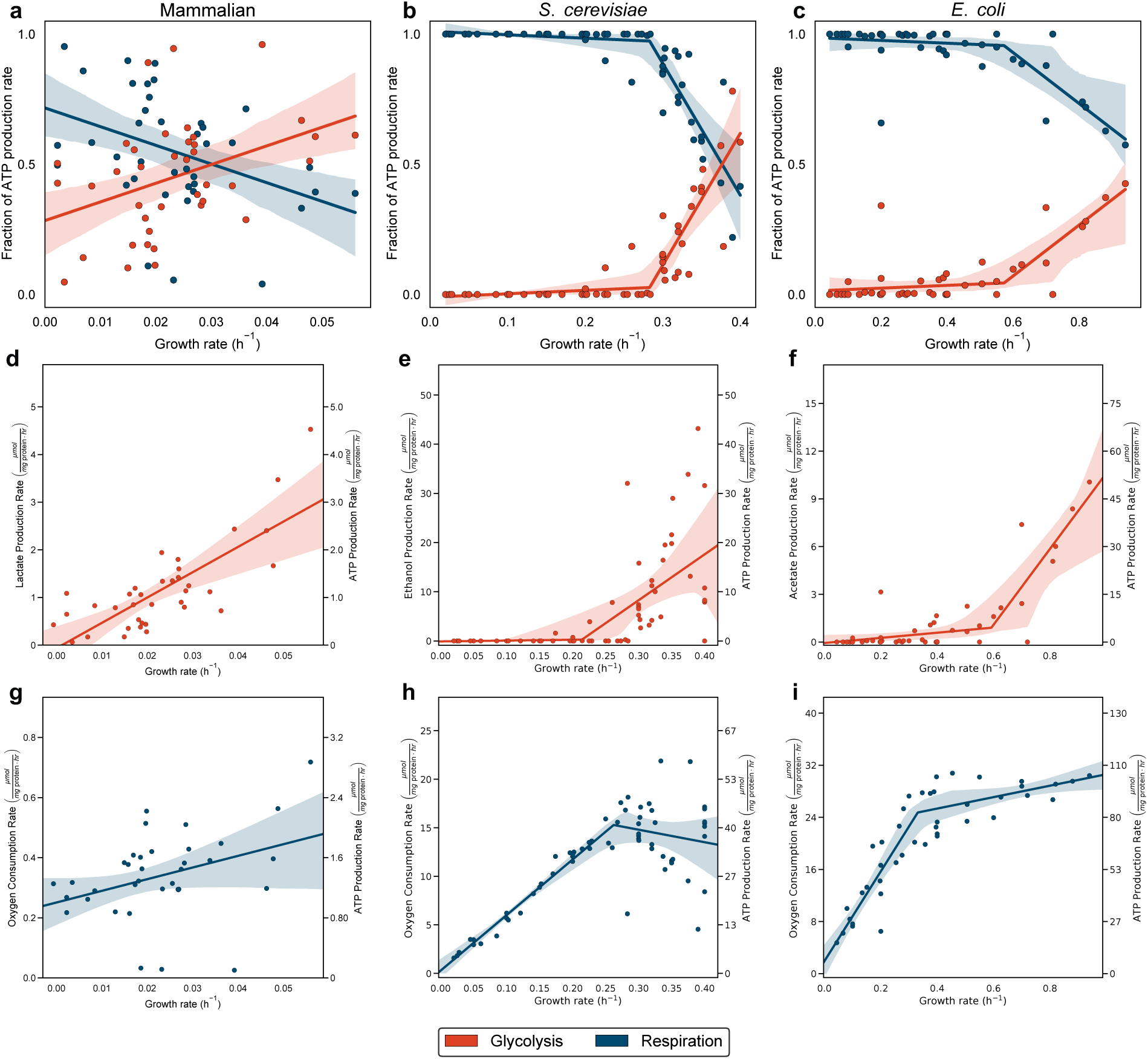
The glycolytic fraction of ATP production increases with growth rate across species. (**a–c**) Fraction of total ATP production derived from glycolysis (red) and respiration (blue) as a function of growth rate for mammalian cells (**a**), *S. cerevisiae* (**b**), and *E. coli* (**c**). (**d–f**) Fermentation product secretion rate versus growth rate; lactate in mammalian cells (**d**), ethanol in *S. cerevisiae* (**e**), and acetate in *E. coli* (**f**). (**g–i**) Oxygen consumption rate versus growth rate for mammalian cells (**g**), *S. cerevisiae* (**h**), and *E. coli* (**i**). In **d–i**, left axes show measured metabolic rates and right axes show the corresponding ATP production rate calculated from organism-specific stoichiometry. Each point represents a single condition or cell line. Solid lines show piecewise linear or linear fits selected by a three-gate criterion (F-test, segment-structure test, and >2-fold slope-change); shaded bands show 95% bootstrap confidence intervals from 10,000 resamples computed on the selected model.

The increased reliance on less efficient glycolysis is conserved across the nearly three billion years of evolutionary distance separating *E. coli*, *S. cerevisiae*, and mammalian cells^53^. This conserved pattern across organisms with fundamentally different cell architectures and metabolic organizations suggests that the shift toward glycolysis at fast growth rates is not a cell-type-specific adaptation but a universal consequence of the kinetic properties of ATP-producing pathways.

### Glycolysis is required to sustain the fastest growth rates observed in microbes and mammalian cells

We next quantified the growth-rate advantage conferred by reliance on glycolytic ATP production for rapid growth, resulting from the Warburg Effect. To estimate the maximum growth rate supportable by each pathway, we can rewrite Eq (1) to get

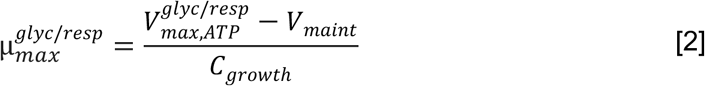

where *μ_max_^glyc/resp^* is maximal growth rate achieved in the presence of only respiration or only glycolysis and *V_max,ATP_^glyc/resp^* is maximal cellular ATP production rate achieved in the presence of only respiration or only glycolysis. The latter values assume that the entire ATP-producing proteome is devoted exclusively to glycolysis or respiration, a theoretical upper bound that no cell achieves, but one that defines the absolute capacity limit of each pathway. Thus, all the values except *μ_max_^glyc/resp^* are estimated from experiments. The estimated maximal ATP production rates (see *Method S2: Detailed calculation of maximal ATP production capacity* for details of the calculation) via respiration are 2.7, 47, and 90 µmol per mg protein per hour for mammalian cells, *S. cerevisiae*, and *E. coli*, and via glycolysis 8.5, 95, and 200 µmol per mg protein per hour, reflecting a consistent 2–3-fold advantage of glycolytic over respiratory capacity across all three organisms. Using Eq. (2), we get that respiration alone can sustain maximum growth rates of 0.024 hr⁻¹ for mammalian cells, 0.29 hr⁻¹ for *S. cerevisiae*, and 0.56 hr⁻¹ for *E. coli*, corresponding to 43%, 73%, and 60% of the fastest observed growth rates in each dataset, respectively (Fig. 4). Glycolytic capacity, by contrast, is sufficient to support the fastest observed growth rates in all three organisms. In mammalian cells, the respiratory capacity limit corresponds to a doubling time of approximately 30 hours. Consistent with this prediction, cells forced to rely predominantly on respiratory ATP production cannot sustain doubling times faster than approximately 30 hours. HeLa cells on galactose double every 37 hours^54^, ovarian cancer cells in glucose-free medium slow to a 40-hour doubling time^55^, LNCaP prostate cancer cells under glucose deprivation slow to approximately 165 hours, and DU145 cells are unable to sustain growth^56^, LDHA/LDHB double-knockout cells slow to 23–31 hours^57^, and pancreatic β-cell lines, which silence LDHA as part of a disallowed gene program, are characteristically slow-cycling with doubling times of 40 hours to several days^58,59^ (Supplementary Text 1). Conversely, cells lacking functional mitochondria sustain near-normal proliferation when provided an alternative means of NAD⁺ regeneration^60–64^, further supporting that glycolytic rather than respiratory ATP production is the limiting factor for rapid growth.

**Fig. 4.**
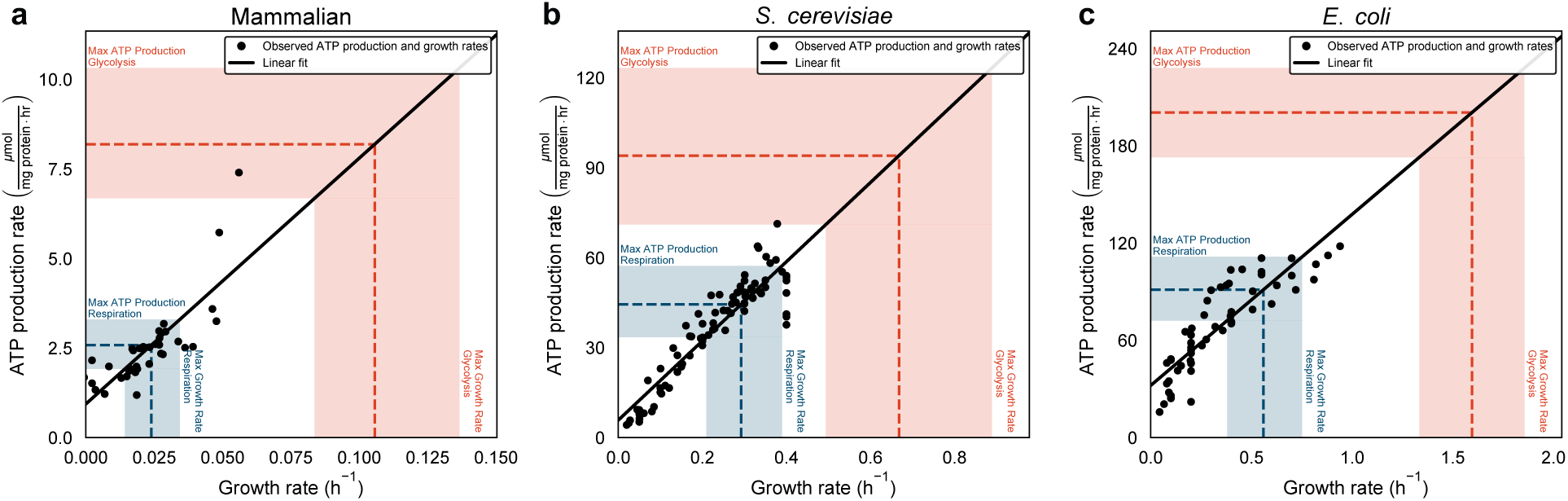
Glycolysis is required to sustain growth rates above the respiratory capacity limit. (a–c) ATP production rate versus growth rate for mammalian cells (a), *S. cerevisiae* (b), and *E. coli* (c), with horizontal dashed lines indicating estimated maximal ATP production capacities via respiration (blue) and glycolysis (red), assuming complete allocation of the ATP-producing proteome to a single pathway. Vertical dashed lines and shaded bands indicate the maximal growth rates (mean ± 95% CI) supportable by each pathway. Shaded bands on capacity estimates represent 95% CIs from 10,000 bootstrap resamples.

## Discussion

Our data provide rigorous estimates of growth-associated and maintenance-associated ATP demands in mammalian cells. Across a diverse panel of twelve cell lines, ATP production scales linearly with growth rate, revealing a conserved energetic cost of biomass accumulation that closely mirrors values previously reported in *E. coli* and *S. cerevisiae*. In rapidly proliferating mammalian cells, growth-associated costs dominate total ATP expenditure, whereas maintenance costs represent a comparatively smaller fraction of cellular energy use, directly contradicting prior reports^34,37,38^. Notably, the inferred maintenance demand in mammalian cells is similar in magnitude to basal metabolic rates estimated for human tissues, consistent with the ability of slow-growing or non-proliferating cells to rely on respiration alone^65^. These results indicate that while maintenance energy requirements are relatively constant, increases in proliferation impose a substantial and conserved ATP burden driven by biosynthetic demand. A key reason microbes achieve faster growth rates than mammalian cells is their ability to generate substantially more ATP per unit biomass^25^. Maximal glycolytic ATP production is 4-to 6-fold higher and respiratory ATP production is 8-to 10-fold higher in *S. cerevisiae* and *E. coli* than in mammalian cells, and microbes allocate a larger fraction of their proteome to ATP-producing enzymes^25^. Because the ATP cost of biosynthesis per gram of new cell mass is broadly conserved across species, these kinetic and allocation differences directly constrain the maximum growth rate achievable by mammalian cells, and suggest that the higher ATP production rate per gram of pathway protein is a highly evolvable trait that has been strongly optimized in microbes (Supplementary Text 2).

Our results support the hypothesis that the high ATP demands of fast growth cannot be satisfied by respiration alone in *E. coli*, *S. cerevisiae*, or mammalian cells, even if the entire ATP-producing proteome were devoted to that pathway (Fig. 3). Glycolytic ATP production is therefore necessary for achieving fast growth rates in all three organisms. Using pharmacological inhibitors of glycolysis and respiration, we directly tested this premise and found that loss of glycolysis could not be compensated by respiration, whereas loss of respiration was fully compensated by glycolysis, sustaining growth at nearly unchanged rates (Fig. 2). Genetic ablation of fermentative glycolysis via LDHA/LDHB double knockout reduces proliferation rate in LS174T and B16 cells^57^, supporting that oxidative phosphorylation alone cannot support maximal proliferative output. We also observed impaired growth upon enforced upregulation of ATP consumption using ATPGobble, providing direct evidence that the capacity for cellular energy production can constrain proliferative potential. This finding addresses a longstanding debate in the cancer metabolism field regarding whether ATP supply poses a genuine barrier to cell growth^34,35^.

Our data provide a quantitative explanation for the Warburg Effect in proliferating cells under nutrient-rich, oxygen-replete conditions, but raise the question of whether the same logic extends to tumors. Tumors derive large fractions of their ATP from aerobic glycolysis^14,66^, even though their net doubling times are far slower than the approximately 30-hour threshold we identify here. Several factors may contribute. First, the slow net doubling time of tumors obscures the actual division rate of individual cells. Thymidine and BrdU labeling studies consistently show that individual tumor cells divide far faster than net tumor growth implies, with cell loss factors of 50–99% accounting for the difference between cell birth rates and volume doubling times^67–69^. Individual cancer cells may therefore be cycling at rates fully consistent with the glycolytic ATP demands our framework predicts. Second, tumor hypoxia drives lactate production independently of the growth-rate-driven mechanism we describe, as substantial tumor fractions experience oxygen tensions insufficient to sustain full electron transport chain flux^70,71^. Third, the lower proteome cost of glycolysis may free proteomic space in cancer cells for purposes other than ATP production for rapid cell growth, such as increasing survival by expressing stress-resistance pathways. Consistent with this, pancreatic tumors have been shown to dramatically downregulate the protein synthesis programs that define the exocrine pancreas, shedding their tissue-specific functions to enable proliferation^72^. Finally, multiple lines of evidence suggest that tumors can grow *in vivo* without glycolytic fermentation, consistent with our hypothesis that it is only required for very rapid growth rates. For example, insulinomas often do not express lactate dehydrogenase but can still form tumors, and xenografts with lactate dehydrogenase deletion^57^ are still capable of forming tumors. How our framework extends to cancer cells *in vivo*, where nutrient availability, hypoxia, and cell loss factors all independently drive lactate production, is a next step toward a quantitative understanding of aerobic glycolysis in tumors. Together, our results establish glycolytic ATP production as a quantitative requirement for rapid mammalian cell growth and provide a mechanistic foundation for understanding the Warburg Effect.

## Methods

### Cell lines and cell culture

All cell lines were cultured in Dulbecco’s modified eagle’s medium [DMEM (Gibco^TM^ 12800082), 3.7 g/L NaHCO_3_, 10% FBS (Gibco^TM^ 10437028), and 100 U / mL Penicillin-Streptomycin (Gibco^TM^ 15140122)]. All experiments were performed in the absence of Penicillin-Streptomycin. The following cell lines were used and obtained from the University of California, Berkeley Cell Culture Facility: C2C12 (ATCC CRL-1772), HeLa (ATCC CCL-2), U-2 OS (ATCC HTB-96), A549 (ATCC CCL-185), SK-BR-3 (ATCC HTB-30), NCI-H1703 (ATCC CRL-5889), PC-3 (ATCC CRL-1435), A431 (ATCC CRL-1555), and MCF-7 (ATCC HTB-22), B16-F10 (ATCC CRL-6475). Huh-7 and MC38 cells were kindly provided by Anders Näär’s and Michael DuPage’s labs, respectively.

### Cell Proliferation

Cell proliferation was quantified by automated nuclei counting on a Cytation 1 imaging reader (Gen5 3.05 software). Cells were seeded in black, clear-bottom 96-well plates (Costar 3904) at densities optimized per line (300–500 cells per well) in 200 µL complete medium. Plates were fixed daily for up to 7 days in 4% paraformaldehyde (diluted from 16% stock in PBS, filtered through a 0.22 µm membrane) for 15 min at 37°C, washed, and stained with Hoechst nuclear dye (1 µg/mL final, from 10,000× stock in PBS). Plates were sealed with light-blocking film and stored at 4°C until imaging. Nuclei were identified by automated segmentation with object size, rolling ball diameter, and thresholding parameters optimized per line. Outer wells were excluded to reduce edge effects, and wells with fewer than 100 nuclei were excluded from analysis. Growth rate was determined by fitting cell number to an exponential growth model:

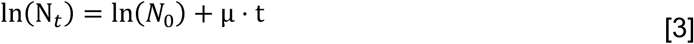

where N₀ and N_t_ are nuclei counts at time 0 and time t (days), and µ is the exponential growth rate constant. Time points after cultures reached confluence were excluded. Rates were converted to hr⁻¹ by dividing µ by 24. Growth metrics were calculated from replicate-level regression fits rather than averaged growth curves (Supplementary Data).

### Measurement of lactate production rate

Lactate production rate was measured using the L-Lactate Assay Kit-I (Eton Bioscience 120001400A). Cells were seeded at the volumetric equivalent of 500,000 HeLa cells (mean HeLa cell volume 2,300 fL) per well of a 6-well plate in 3 mL DMEM and incubated at 37°C in 5% CO₂. After 24 hr, medium was replaced with 3 mL assay medium (DMEM, Gibco 12100061; 10% dialyzed FBS, Gibco 26400044; 3.7 g/L NaHCO₃). Media samples of 200 µL were collected hourly for 4 hr and immediately frozen on dry ice, then stored at −80°C. At the end of the sampling period, cells were trypsinized and counted by Coulter counter. For analysis, 50 µL of each sample was combined with 50 µL L-Lactate assay reagent and absorbance at 490 nm was measured for 45 min at 37°C on a Cytation 1. Lactate concentrations were determined from a standard curve of OD₄₉₀ slope versus known L-lactate concentration and converted to moles per mg protein using cell counts and per-cell protein concentrations. Lactate production rate was determined by linear regression across the four-hour time course (Supplementary Data).

### Measurement of Oxygen Consumption Rate

Oxygen consumption rate (OCR) was measured using the Agilent Seahorse XFe24 Analyzer. Cells were seeded at the volumetric equivalent of 100,000 HeLa cells per well of an XFe24 microplate in 150 µL DMEM, where mean HeLa cell volume was determined to be 2,300 fL by Coulter counter. After 24 hr, medium was replaced with 500 µL assay medium (DMEM, Gibco 12100061; 10% dialyzed FBS, Gibco 10437028; 5 mM HEPES-KOH, pH 7.4) and plates were transferred to the analyzer. Each measurement cycle consisted of a 2 min mix, 2 min wait, and 4 min measurement period. Three basal measurements were collected, followed by sequential injections of oligomycin (1 µM final) and antimycin A with rotenone (1 µM each final), with three measurements collected after each injection. All compounds were delivered as 50 µL concentrated solutions in assay medium. Following the assay, cells were trypsinized and counted on a Coulter Z2 Counter; OCR values were normalized to mg protein per well using per-cell protein concentrations derived from measured cell volumes.

### Cell volume and protein content

Cell volume and concentration were measured at during cell passaging and under all experimental conditions using a Coulter Z2 Counter Cell Particle Analyzer (Beckman). Mean cell volume (fL) and standard deviation were determined from repeated measurements for each cell line and used throughout the study to convert cell counts to protein mass (Supplementary Data). Protein concentration was determined by BCA assay (Pierce 23225). One million cells were pelleted at 600 × g, washed twice with PBS to remove media protein, and lysed in 500 µL of 1% SDS at 90°C for 10 min. Samples were assayed in triplicate against a bovine serum albumin standard curve with absorbance measured at 562 nm on a Cytation 1.

### ATP production rate calculation

Total cellular ATP production rate was calculated as:

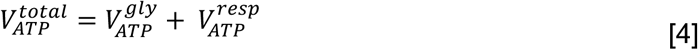

where all rates are in µmol mg protein⁻¹ hr⁻¹. Glycolytic ATP production was calculated from measured byproduct secretion rates, acetate, ethanol, and lactate for *E. coli*, *S. cerevisiae*, and mammalian cells, respectively, at yields of 5, 1, and 1 ATP per byproduct molecule^25^. Respiratory ATP production was calculated from OCR using stoichiometric ATP yields per glucose of 20, 16, and 24 for *E. coli*, *S. cerevisiae*, and mammalian cells, respectively, derived from experimental measurements and stoichiometric accounting of proton translocation, proton leak, and ATP synthase H⁺/ATP ratios as described previously^25^. For mammalian cells, non-mitochondrial OCR was measured as the residual OCR after antimycin A and rotenone treatment and subtracted from basal OCR before calculating respiratory ATP. These calculations assume glucose as the respiratory substrate; the P/O ratio varies by less than 20% across physiologically relevant substrates, so this assumption does not affect our conclusions.

### Maximal ATP production capacity

Maximal ATP production rates via glycolysis and respiration were estimated by combining three parameters: the maximal specific activity of each pathway (*V_max_^pathway^*, µmol glucose mg pathway protein⁻¹ min⁻¹), the ATP yield per glucose (γ*_pathway_*), and the fraction of the total proteome allocated to ATP-producing enzymes (*ϕ_total_^ATP^*), as

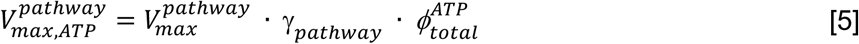

For *E. coli* and *S. cerevisiae*, *V_max_^pathway^* for glycolysis was estimated from maximal acetate and ethanol production rates in batch culture; *V_max_^pathway^* for respiration was estimated from maximal OCR on non-fermentable substrate. For mammalian cells, *V_max_^pathway^* for glycolysis was estimated from oligomycin-maximized lactate production rates, and *V_max_^pathway^* for respiration from FCCP-stimulated OCR after subtraction of antimycin A-insensitive non-mitochondrial respiration. Proteome fractions for mammalian cells were derived from published quantitative proteomics datasets for 11 cell lines of distinct tissue origins (see Method S4. Estimation of proteome fractions allocated to ATP-producing enzymes). Uncertainty was propagated by nonparametric bootstrap resampling over 10,000 iterations; 95% CIs are the 2.5th–97.5th percentiles of the resulting distributions.

### Pharmacological perturbations

LDH was inhibited with GSK-2837808A (0, 50, 100, 150 µM; Cayman Chemical). Complex III was inhibited with antimycin A (1 µM; Sigma). Translation was inhibited with cycloheximide (0, 0.05, 0.2, 1 µM; Sigma). For each perturbation, growth rate, lactate production rate, and OCR were measured simultaneously in MCF-7, U-2 OS, and C2C12 cells as described above.

### Production of cells stable ATPGobble under control of doxycycline promoter

Doxycycline-inducible pLVX-TetOne vectors encoding codon-optimized *E. coli* F₁-ATPase subunits *atpA*, *atpG*, and *atpD* (Thermo Fisher codon optimizer; backbone pLVX-TetOne-Puro, Takara 631849) were used to express ATPGobble. The puromycin resistance cassette in the *atpA* and *atpD* vectors was replaced with blasticidin (Addgene 183751) and zeocin (Addgene 161748) resistance, respectively. A catalytically inactive control was generated by introducing an *atpD*(K155Q) mutation by site-directed mutagenesis (NEB E0554S). Lentivirus was produced in HEK293T cells seeded at 5 × 10⁵ cells per well of a 6-well plate in 2 mL high-glucose DMEM (Life Technologies 11995) with 10% FBS. After 24 hr, medium was replaced and cells were transfected with 500 ng psPAX2 (Addgene 12260), 50 ng pMD2.G (Addgene 12259), and 500 ng transfer vector, combined in 50 µL Opti-MEM (Life Technologies 31985-070) and mixed with 3 µL X-tremeGENE 9 (Roche 06365787001) diluted in 50 µL Opti-MEM, incubated 30 min at room temperature, then added to cells. Virus-containing supernatant was harvested 48 hr post-transfection, centrifuged at 500 × g for 5 min, and filtered through a 0.45 µm PES membrane (Millipore SLHP033RS). Target cells were seeded at 50,000 cells per well in 2 mL growth medium 24 hr before infection. Polybrene was added to 8 µg/mL and 1 mL viral supernatant per construct was applied. Medium was replaced 24 hr post-infection, and selection was initiated at 48 hr with blasticidin (5 µg/mL), zeocin (100 ng/mL), puromycin (10 µg/mL), and geneticin (500 µg/mL). Cultures were maintained under selection until uninfected control wells were fully eliminated. ATPGobble was induced with doxycycline (1 µg/mL) for 72 hr before measurement.

### Biosynthetic ATP cost estimates

The ATP required to support cellular biomass synthesis was estimated by summing contributions from protein, RNA, DNA, and lipid synthesis, weighted by measured biomass compositions, as

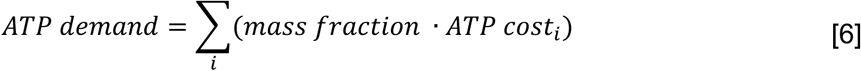

Protein synthesis costs approximately 5.0 ATP equivalents per amino acid for mammalian cells (4 ATP for tRNA charging and ribosomal elongation, plus 0.40 ATP for amino acid import via Na⁺-coupled transporters) and 5.7 ATP equivalents per amino acid for microorganisms grown in minimal medium (including de novo amino acid biosynthesis at 1.4 ATP per amino acid). RNA synthesis costs approximately 10 ATP equivalents per ribonucleotide for mammalian cells and 12 for microorganisms, reflecting the cost of de novo nucleotide biosynthesis. DNA synthesis costs approximately 10 and 12 ATP equivalents per deoxyribonucleotide for mammalian and microbial cells, respectively, including additional replication costs. Lipid synthesis costs approximately 16 ATP equivalents per phospholipid for mammalian cells under serum-replete conditions (reflecting a 70%/30% exogenous/*de novo* split) and 22–39 ATP equivalents for microbial cells. Full details are provided in Method S3 Detailed biosynthetic ATP cost calculations.

### Statistics

All measurements are reported as mean ± 95% CI from 10,000 bootstrap resamples unless otherwise noted. Linear regressions were performed by ordinary least squares. Piecewise linear models with a single breakpoint were fit by nonlinear least squares with the breakpoint constrained to lie within the interior of the data range. Model selection between linear and piecewise fits required three criteria to be met, otherwise the linear model was retained: (1) an F-test favoring piecewise over linear (p < 0.05); (2) statistically meaningful trends within each segment; and (3) a greater than 2-fold change in slope between segments, with sign reversals automatically satisfying this criterion. Bootstrap 95% confidence intervals on the fitted curves were computed conditional on the selected model. Model selection was performed once on the full dataset and all 10,000 resamples were forced to fit the same model family. R² was computed as 1 − SS_res / SS_tot, and p-values were obtained from an F-test against an intercept-only null. All analyses were performed in Python 3.10 using NumPy and SciPy.

## Data availability

All data are available in the main text or the supplementary data.

## Code availability

All data and code used in the figure generation are available as a GitHub repository via https://github.com/DenisTitovLab/ATPGrowth.

## Acknowledgments

Research reported in this publication was supported by the National Institute of General Medical Sciences (NIGMS) of the National Institutes of Health (NIH) under award number R35GM152114 to D.V.T. and by the Cancer Research Coordinating Committee of the University of California under Faculty Seed Award Grant Number C24CR7706 (D.V.T.) and Predoctoral Fellowship (M.A.K.). We gratefully acknowledge the University of California, Berkeley Cell Culture Facility for assistance with cell culture and maintenance.

## Author contributions

M.A.K. and D.V.T. designed research; M.A.K., S.Z., and B.T.H. performed experiments; M.A.K. and A.E. contributed new reagents/analytic tools; M.A.K. and D.V.T. analyzed data; and M.A.K. and D.V.T. wrote the paper

## Supplemental Information

### Contents

- Figure S1: Oxygen consumption rate, lactate production rate, and growth rate of twelve mammalian cells lines
- Figure S2: Comparison of ATP cost of biomass production across organisms
- Figure S3: Effects of glycolysis inhibition, respiration inhibition, biosynthesis inhibition, and ATPGobble expression on growth rate, oxygen consumption, and lactate production.
- Supplementary Text 1: Supporting evidence for the approximately 30-hour respiratory capacity limit
- Supplementary Text 2: Species comparison of ATP production capacity
- Method S1: Detailed calculation of cellular ATP production rates from glycolysis and respiration
- Method S2: Detailed calculation of maximal ATP production capacity
- Method S3: Detailed biosynthetic ATP cost calculations
- Method S4: Estimation of proteome fractions allocated to ATP-producing enzymes
- Method S5: Conversion of published values to common units

**Supplementary Text 1: Supporting evidence for the approximately 30-hour respiratory capacity limit**

Our calculations predict that mammalian cells relying exclusively on respiration cannot sustain doubling times faster than approximately 30 hours. This threshold is determined by the intersection of the estimated maximal respiratory ATP production rate (2.5 µmol ATP mg protein⁻¹ hr⁻¹; 95% CI: 1.9–3.3) with the linear ATP demand–growth relationship from Fig. 1a. We note that this threshold is a model-derived estimate with compounding uncertainties from the proteome fraction, maximal specific activity, and P/O ratio measurements; the 95% CI on the threshold spans approximately 25–38 hours.

Several independent lines of experimental evidence are consistent with this prediction.

### Galactose substitution

HeLa cells grown on galactose rather than glucose are forced to rely predominantly on oxidative phosphorylation for ATP because galactose enters glycolysis at a rate insufficient to support significant lactate production. Under these conditions, HeLa cells double every 37 hours, compared to approximately 24 hours on glucose^1^. This 37-hour doubling time falls above the predicted approximately 30-hour threshold, consistent with the prediction.

### LDHA/LDHB double knockout

Genetic ablation of fermentative lactate production through double knockout of LDHA and LDHB forces complete reliance on oxidative phosphorylation for NAD⁺ regeneration and ATP production. This intervention reduces proliferation to doubling times of 31 hours in human LS174T colon adenocarcinoma cells and 23 hours in murine B16 melanoma cells^2^. Both values are at or above the predicted threshold, and the B16 value (23 hours) falls within the lower bound of the 95% CI (approximately 25 hours), consistent with the prediction.

### Cancer cells under glucose deprivation

Ovarian cancer cells in glucose-free medium slow to a 40-hour doubling time^3^. LNCaP cells under glucose-free conditions slow to a doubling time of approximately 165 hours, and DU145 cells cannot sustain growth^4^. While glucose deprivation affects multiple metabolic processes beyond glycolytic ATP production, these observations are consistent with the conclusion that cells unable to maintain glycolytic ATP flux cannot sustain rapid proliferation.

### Pancreatic beta cell lines

Pancreatic beta cells silence LDHA expression as part of a beta-cell-specific gene program that suppresses aerobic glycolysis. Beta-cell-derived lines are characteristically slow-cycling, with doubling times ranging from approximately 40 hours to several days depending on the line and conditions^5,6^. This is consistent with the prediction that cells unable to use glycolytic ATP production are constrained to doubling times above approximately 30 hours.

### Cells lacking functional mitochondria

Conversely, cells completely lacking functional mitochondria (rho-zero cells) can sustain near-normal proliferation when provided an exogenous means of NAD⁺ regeneration, such as pyruvate supplementation or exogenous aspartate^7–11^. This confirms that glycolytic ATP production alone is sufficient to meet the demands of rapid growth when provided an alternative means of NAD⁺ regeneration.

Taken together, these observations from five independent experimental systems are consistent with the approximately 30-hour threshold predicted by our model, and support the conclusion that glycolytic ATP production is required for doubling times faster than approximately 30 hours.

**Supplementary Text 2: Species comparison of ATP production capacity**

A key quantitative finding of this study is that mammalian cells operate near their respiratory capacity limit even at moderate growth rates, whereas *E. coli* and *S. cerevisiae* have substantial respiratory headroom at slow growth rates and only switch to glycolysis at faster growth rates. This difference arises from two factors: (1) mammalian respiration produces ATP at a lower rate per milligram of pathway protein than microbial respiration, and (2) mammalian cells allocate a smaller fraction of their proteome to ATP-producing enzymes.

As reported in our previous study^12^, the maximal specific activity of the respiratory pathway is approximately 8-to 10-fold lower in mammalian cells than in *E. coli* or *S. cerevisiae*. The maximal specific activity of the glycolytic pathway is approximately 4-to 6-fold lower. Combined with proteome fractions of approximately 5–8% for ATP-producing enzymes in mammalian cells versus 10–20% in microbes, these differences result in maximal ATP production rates that are an order of magnitude lower per unit biomass in mammalian cells.

Because the ATP cost of biosynthesis per gram of new cell mass is broadly conserved across species (reflecting the shared biochemistry of peptide elongation, nucleic acid synthesis, and lipid assembly), the lower ATP production capacity of mammalian cells directly constrains their maximum growth rate. The fastest-growing mammalian cell lines in our dataset (doubling time approximately 14–16 hours) are already operating near the estimated glycolytic capacity limit, whereas the fastest-growing *E. coli* and *S. cerevisiae* strains have substantial additional glycolytic capacity.

This analysis suggests that the maximum growth rate of mammalian cells is constrained primarily by the kinetic properties of their ATP-producing enzymes rather than by the availability of biosynthetic precursors or the cost of biosynthesis itself. Whether the lower specific activity of mammalian respiratory complexes reflects an evolutionary trade-off or simply a lack of selective pressure for faster growth in multicellular organisms remains an open question.

**Method S1. Detailed calculation of cellular ATP production rates from glycolysis and respiration**

Total cellular ATP production rate was calculated as the sum of glycolytic and respiratory contributions, where all rates are in units of µmol mg protein⁻¹ hr⁻¹:

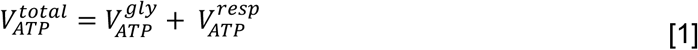

Glycolytic ATP production was calculated from the measured rate of fermentative byproduct secretion at organism-specific stoichiometric yields. Respiratory ATP production was calculated from the measured oxygen consumption rate using stoichiometric ATP yields per glucose derived from experimental measurements combined with accounting of proton translocation efficiency, proton leak, and ATP synthase H⁺/ATP ratios^12^.

### E. coli

The primary glycolytic byproduct in *E. coli* is acetate, produced via the Pta-AckA pathway. This pathway converts glucose to acetate with a net yield of 10 ATP per glucose (2 ATP from substrate-level phosphorylation and 8 ATP from ETC oxidation of 4 NADH produced per glucose). Because the NADH oxidation step consumes oxygen, this respiratory contribution is already captured in the measured OCR. To avoid double-counting, the oxygen consumed by Pta-AckA-linked NADH oxidation was subtracted from total OCR before calculating respiratory ATP. The organism-specific equation is therefore:

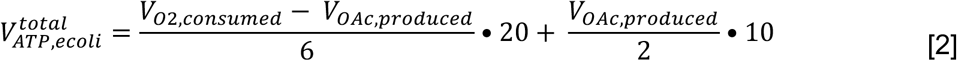

where *V_o2,consumed_* and *V_OAC,produced_* are in µmol mg protein⁻¹ hr⁻¹, the factor of 6 converts O_2_ consumed per glucose to moles of glucose fully oxidized, and 20 is the ATP yield per glucose from full respiratory oxidation in *E. coli*.

### S. cerevisiae

Ethanol is the primary fermentative byproduct in *S. cerevisiae* (2 mol ethanol per mol glucose, yielding 2 ATP), and oxygen consumption reflects full glucose oxidation through respiration. The organism-specific equation is:

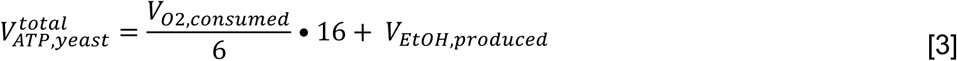

where 16 is the ATP yield per glucose from full respiratory oxidation in *S. cerevisiae*.

### Mammalian cells

Lactate is the primary fermentative byproduct in mammalian cells (2 mol lactate per mol glucose, yielding 2 ATP by substrate-level phosphorylation). A fraction of total OCR is attributable to non-mitochondrial processes that do not contribute to ATP production. This non-mitochondrial OCR was measured as the residual OCR remaining after treatment with antimycin A and rotenone, which together abolish mitochondrial electron transport, and was subtracted from basal OCR before calculating respiratory ATP:

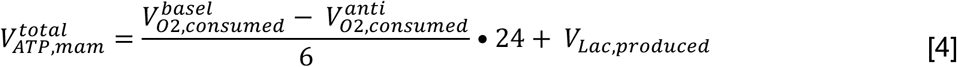

where 24 is the ATP yield per glucose from full respiratory oxidation in mammalian cells.

These calculations assume glucose as the sole respiratory substrate for the purpose of assigning a single P/O ratio. In practice, mammalian cells in standard culture conditions oxidize a mixture of substrates, primarily glucose, glutamine, and fatty acids. The P/O ratio differs by less than 20% across these substrates: glucose oxidation yields approximately 4.0 ATP per O2, glutamine oxidation approximately 3.7 ATP per O2, and fatty acid oxidation approximately 3.5 ATP per O2. If mammalian cells rely more heavily on glutamine or fatty acid oxidation, as is commonly observed in standard serum-replete culture conditions^1,13^, the effective P/O ratio would be modestly lower than the glucose-based value used here, meaning our calculations slightly overestimate respiratory ATP production. This would only strengthen the central conclusion that respiration alone is insufficient to sustain rapid cell growth.

### Method S2. Detailed calculation of maximal ATP production capacity

The maximal ATP production rate achievable if the entire ATP-producing proteome were devoted exclusively to a single pathway was calculated as:

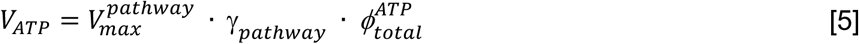

where *V_max_^pathway^* is the maximal specific activity of the pathway (µmol glucose mg pathway protein⁻¹ min⁻¹), γ*_pathway_* is the ATP yield per glucose consumed (ATP per glucose), and *ϕ_total_^ATP^* is the fraction of the total cellular proteome allocated to ATP-producing enzymes.

*V_max_^pathway^* is defined as the µmol of substrate consumed per minute per milligram of pathway protein, analogous to the specific activity of individual enzymes and equivalent to what others have termed proteome efficiency. For *E. coli* and *S. cerevisiae*, *V_max_^pathway^* for glycolysis was estimated from maximal acetate and ethanol production rates, respectively, measured in batch culture on a fermentable carbon source at the point of maximal fermentative flux. *V_max_^pathway^* for respiration was estimated from maximal oxygen consumption rates measured in batch culture or accelerostat on a non-fermentable carbon source. For mammalian cells, *V_max_^pathway^* for glycolysis was estimated from oligomycin-maximized lactate production rates, which block mitochondrial ATP synthesis and thereby drive maximal glycolytic flux. *V_max_^pathway^* for respiration was estimated as the difference between FCCP-stimulated OCR and antimycin A-insensitive non-mitochondrial OCR, isolating maximal mitochondrial oxygen consumption capacity. All values are reported in Supplementary Data.

γ*_pathway_* values are identical to those used in the basal ATP production rate calculations described in Supplementary Method 1. For glycolysis, the yield is 2 ATP per glucose for *S. cerevisiae* and mammalian cells and 10 ATP per glucose for *E. coli*. For respiration, the yield is 20, 16, and 24 ATP per glucose for *E. coli*, *S. cerevisiae*, and mammalian cells, respectively. The same caveats regarding respiratory substrate composition apply.

*ϕ_total_^ATP^* represents the fraction of total cellular protein mass attributable to glycolytic or respiratory enzymes, estimated using the total protein approach applied to published quantitative mass spectrometry proteomics datasets. For *E. coli* and *S. cerevisiae*, proteome fractions were derived from published quantitative proteomics datasets as described previously^12^. For mammalian cells, proteome fractions were derived from 11 cell lines of distinct tissue origins (A549, GAMG, HeLa, HepG2, Jurkat, K562, LNCaP, MCF7, RKO, U-2 OS, and HEK293). All values are reported in Supplementary Data.

To account for experimental variability in enzyme abundance, pathway allocation, and substrate uptake rates, uncertainty was propagated by nonparametric bootstrap resampling over 10,000 iterations. In each iteration, glucose uptake rates for fermentation and respiration and total proteome allocations to ATP-producing enzymes were independently sampled with replacement from their respective empirical distributions. *V_ATP_* was calculated for each bootstrap replicate. After 10,000 iterations, the 2.5th and 97.5th percentiles of the resulting distribution were taken as the 95% confidence interval for maximal ATP production capacity via each pathway. This procedure propagates uncertainty from all three input parameters simultaneously and does not assume any particular parametric distribution for the inputs.

### Method S3. Detailed biosynthetic ATP cost calculations

The ATP required to support cellular biomass synthesis was estimated by summing contributions from each major macromolecular class, weighted by measured biomass compositions:

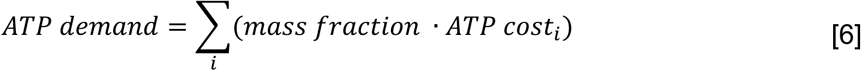

where i ∈ {protein, RNA, DNA, lipid, other}.

Biomass compositions were taken from the literature: for *E. coli*^14–16^, protein 54%, lipid 9.3%, DNA 3.1%, RNA 18%, other 15.6%; for *S. cerevisiae*^17–19^, protein 43%, lipid 7.3%, DNA 2.0%, RNA 5.0%, other 42.7%; for mammalian cells^20–22^, protein 69%, lipid 13%, DNA 1.3%, RNA 4.5%, other 12.2%.

Throughout these calculations, “ATP equivalents” refers collectively to all high-energy phosphate currencies (ATP, GTP, UTP, CTP), which are freely interconverted by nucleoside diphosphate kinase. NADH and NADPH are excluded from this accounting because reducing equivalents do not directly contribute to the measured quantities (fermentation byproduct release and oxygen consumption) used to calculate ATP production rates. Counting reducing equivalents on the demand side while excluding them from the measured supply would introduce a systematic inconsistency in the energy balance.

### Protein synthesis

The total energetic cost of protein synthesis is approximately 4 ATP equivalents per amino acid: 1 ATP for aminoacyl-tRNA synthetase charging, 2 GTP for ribosomal translocation and peptide bond formation, plus overhead for initiation (approximately 1 GTP per ribosome per initiation event, amortized over chain length) and termination^14,23,24^. Additional costs for mRNA synthesis (approximately 0.20 ATP per amino acid, based on mRNA length and nucleotide synthesis costs), ribosomal proofreading (approximately 0.10 ATP per amino acid), and post-translational modification and assembly (approximately 0.006 ATP per amino acid) were included as additive terms^25^.

For *E. coli* and *S. cerevisiae* grown in minimal media, amino acids must be synthesized *de novo*. The average biosynthetic cost was calculated as the weighted mean of the ATP costs of synthesizing each of the 20 amino acids from central metabolic precursors, weighted by the amino acid composition of cellular protein. This yields an average of approximately 1.4 ATP equivalents per amino acid for both organisms^25^. Total protein synthesis cost: approximately 5.7 ATP equivalents per amino acid for microbial cells.

For mammalian cells cultured in amino acid-replete media (DMEM with 10% FBS), amino acids are imported rather than synthesized. The dominant transport systems for amino acids in mammalian cells are Na⁺-coupled symporters (system A, ASC, and N transporters), which co-import one Na⁺ per amino acid^26^. The energetic cost of Na⁺ extrusion by Na⁺/K⁺-ATPase is approximately 1 ATP per 3 Na⁺ ions, yielding approximately 0.33 ATP per amino acid for Na⁺-coupled transport^27^. Accounting for the mixture of Na⁺-coupled and facilitated diffusion transporters, the weighted average import cost is approximately 0.4 ATP per amino acid^26,28^. Total protein synthesis cost: approximately 4.7 ATP equivalents per amino acid for mammalian cells.

### RNA synthesis

*De novo* purine biosynthesis requires 5 ATP equivalents per purine ring (IMP synthesis from PRPP), plus 1 ATP for AMP synthesis or 1 GTP for GMP synthesis, plus 1 ATP for phosphorylation to the triphosphate. De novo pyrimidine biosynthesis requires 2 ATP equivalents per pyrimidine ring (UMP synthesis), plus 1 ATP for phosphorylation to UTP, plus 1 ATP for CTP synthesis from UTP. Weighted by nucleotide abundance in cellular RNA (ATP 26%, GTP 32%, CTP 20%, UTP 22%)^25^, the average cost is approximately 12 ATP equivalents per ribonucleotide for microbial cells. For mammalian cells, glutamine-dependent biosynthetic steps were adjusted to reflect partial import of glutamine from media, reducing the average cost to approximately 10 ATP equivalents per ribonucleotide.

### DNA synthesis

Deoxyribonucleotide synthesis costs were derived from ribonucleotide precursor costs plus the cost of ribonucleotide reductase-mediated reduction (approximately 1 ATP equivalent per nucleotide, via NADPH consumption). Additional replication costs include: helicase-mediated strand separation (approximately 2 ATP per base pair), lagging-strand synthesis and ligation (approximately 1 ATP per nucleotide for Okazaki fragment processing), proofreading and mismatch repair (approximately 0.5 ATP per nucleotide), supercoiling maintenance by topoisomerases (approximately 0.5 ATP per nucleotide), and methylation (approximately 0.1 ATP per nucleotide)^25^. Total: approximately 12 ATP equivalents per deoxyribonucleotide for microbial cells and approximately 10 for mammalian cells.

### Lipid synthesis

The energetic cost of lipid biosynthesis was calculated on a per-phospholipid basis by accounting for fatty-acid synthesis, head-group formation, acyl-chain attachment, and, where applicable, intracellular transport. Fatty-acid composition was assumed to be dominated by C16:0 (43%), C16:1 (33%), and C18:1 (24%) species^25^. Based on established stoichiometry, synthesis of saturated and monounsaturated C16 fatty acids was assigned an energetic cost of 8.0 ATP equivalents per acyl chain, while synthesis of C18 fatty acids was assigned a cost of 9.0 ATP equivalents^29^. Weighting by relative abundance yields an average fatty-acid synthesis cost of 8.2 ATP equivalents per acyl chain. Formation of a glycerophospholipid requires two fatty-acid chains, biosynthesis of the phospholipid head group, and activation and esterification of the acyl chains. Head-group biosynthesis was assigned a cost of 2.0 ATP equivalents per phospholipid, and activation and attachment of two fatty acids to the glycerol backbone was assigned a cost of 4.0 ATP equivalents^30^. Summing fatty-acid synthesis, head-group formation, and acyl-chain attachment yields an average energetic cost of 22 ATP equivalents per phospholipid in prokaryotic cells. In *S. cerevisiae*, additional energetic costs associated with fatty-acid transport into the cytosol from the mitochondria were included. Transport of C16 and C18 fatty acids was assigned costs of 8.0 and 9.0 ATP equivalents, respectively, yielding a weighted average transport cost of 8.2 ATP equivalents per phospholipid. Including this transport term yields a total energetic cost of 39 ATP equivalents per phospholipid in *S. cerevisiae*.

For mammalian cells, the relevant cost depends on the balance between de novo synthesis and scavenging of exogenous fatty acids. Standard culture media supplemented with 10% fetal bovine serum provides BSA-bound free fatty acids and LDL particles, and inhibition of fatty acid synthase has no measurable effect on proliferation under serum-replete conditions, indicating that de novo synthesis is dispensable when exogenous lipids are available. Isotope tracing studies in proliferating mammalian cell lines estimate that approximately 60–70% of cellular lipid carbon is derived from scavenged exogenous fatty acids, while 20–30% is synthesized de novo from glucose^31^. For scavenged fatty acids, the relevant cost is acyl-CoA activation only (2.0 ATP equivalents per chain, or 4.0 ATP equivalents for two chains per phospholipid), plus head-group biosynthesis (2.0 ATP equivalents), totaling 6.0 ATP equivalents per phospholipid^30^. For de novo synthesized fatty acids, the full synthesis plus citrate shuttle cost applies, totaling 39 ATP equivalents per phospholipid as in *S. cerevisiae*. Weighting these two costs by their estimated contributions yields an average cost of 0.7 × 6 + 0.3 × 39 ≈ 16 ATP equivalents per phospholipid for mammalian cells under standard serum-replete culture conditions.

### Other biomass components

In *all three organisms*, a non-negligible fraction of the ATP cost of biomass synthesis arises from macromolecular components outside of protein, RNA, DNA, and bulk lipids. In *E. coli*, include lipopolysaccharide (LPS) components, peptidoglycan precursors, storage carbohydrates, polyamines, and one-carbon metabolism intermediates. Using published biomass compositions, we quantified the abundance of each building block and assigned ATP costs associated with their biosynthesis or activation, such as nucleotide-sugar formation, fatty-acid synthesis, and precursor activation^25^. While the ATP cost of individual components is modest, their cumulative contribution amounts to 1.8 umol ATP per mg protein. In *S. cerevisiae*, the cell wall is composed predominantly of β-1,3-and β-1,6-glucans, mannoproteins, and a smaller fraction of chitin, with polysaccharides accounting for the majority of wall mass^32,33^. Biosynthesis of these polymers proceeds via activated sugar nucleotides (e.g., UDP-glucose, GDP-mannose, UDP-GlcNAc), such that incorporation of each hexose or amino-hexose residue incurs an energetic cost of approximately one ATP equivalent for nucleotide-sugar activation^34^. In addition to structural polysaccharides, trehalose is present in actively growing yeast cells as a minor storage carbohydrate. Quantitative measurements in continuous (chemostat) cultures report trehalose levels of 33 mg gCDW⁻¹ (3.3% dry weight), with trehalose content decreasing as growth rate increases, consistent with a limited reserve role during exponential growth^35^. These values support inclusion of both cell wall polysaccharides and trehalose as modest but non-negligible contributors to the ATP cost of biomass synthesis in growing yeast. In mammalian cells, the corresponding “other” biomass fraction differs from microbes and is dominated by protein-and lipid-linked glycans, with smaller contributions from sterols, glycogen, and polyamines. Mammalian cells are extensively glycosylated, and N-linked and O-linked glycans and glycosphingolipids together account for 5 percent of cellular dry mass, depending on cell type and growth conditions^36–38^. As in yeast, glycan biosynthesis proceeds through activated sugar nucleotides (e.g., UDP-hexoses and UDP-N-acetylhexosamines), such that incorporation of each sugar residue requires nucleotide-sugar activation and incurs an ATP-equivalent energetic cost^39,40^. In addition, small but measurable pools of glycogen and polyamines contribute further ATP demand in proliferating cells^21,41,42^. Using conservative estimates for these components, we estimate that mammalian “other” biomass components contribute 0.5 µmol ATP per mg protein, a modest but non-negligible fraction of total biomass synthesis cost.

### Method S4. Estimation of proteome fractions allocated to ATP-producing enzymes

The fraction of the total proteome allocated to ATP-producing enzymes (*ϕ_total_^ATP^*) was estimated from published quantitative mass spectrometry datasets.

For mammalian cells, proteome fractions were estimated from 11 cell lines of distinct tissue origins: A549 (lung), GAMG (glioblastoma), HeLa (cervical), HepG2 (hepatocellular), Jurkat (T cell leukemia), K562 (chronic myelogenous leukemia), LNCaP (prostate), MCF-7 (breast), RKO (colon), U-2 OS (osteosarcoma), and HEK293 (embryonic kidney). For each cell line, the mass fraction of glycolytic and respiratory enzymes was calculated from published protein abundance data^43,44^. The total ATP-producing proteome fraction (*ϕ_total_^ATP^*) was defined as the sum of glycolytic and respiratory enzyme fractions (Supplementary Data).

For *E. coli*, proteome fractions were derived from published quantitative proteomics data from cells grown at multiple growth rates in glucose minimal medium. For *S. cerevisiae*, proteome fractions were derived from published data from cells grown in glucose medium (Supplementary Data).

Uncertainty in *ϕ_total_^ATP^* was propagated by bootstrap resampling across cell lines (for mammalian cells) or growth conditions (for microbes), sampling with replacement from the empirical distribution of *ϕ_total_^ATP^* The 2.5th and 97.5th percentiles of the resulting distribution provided 95% confidence intervals.

### Method S5. Conversion of published values to common units

Previously reported ATP requirements were converted to µmol ATP per mg protein for direct comparison with our measurements.

### Microbial datasets

For datasets reported per gram dry weight (gCDW), protein mass fractions of 0.55 g protein per gCDW (*E. coli*) and 0.44 g protein per gCDW (*S. cerevisiae*) were used for conversion.

### Mouse LS cells (Sinclair 1974)

Growth-associated ATP consumption of 22 pmol ATP per cell and maintenance-associated consumption of 19 pmol ATP per cell per day were reported^45^. Protein content was inferred from Fig. 1 of that study as approximately 100 pg protein per cell, yielding 220 µmol ATP per mg protein (growth) and 7.9 µmol ATP per mg protein per hour (maintenance).

### Mouse LS cells (Kilburn et al. 1969)

Growth-associated ATP consumption of 23 pmol ATP per cell and maintenance of 17 pmol ATP per cell per day were reported^46^. A dry weight of 6.6 × 10⁻¹⁰ g per cell and a protein fraction of 69% were used for conversion, yielding 51 µmol ATP per mg protein (growth) and 1.4 µmol ATP per mg protein per hour (maintenance).

### Basal metabolic rate in humans

The standard metabolic equivalent of task (MET) for resting metabolism is 3.5 mL O₂ per kg wet weight per minute^47^. Converting to ATP flux per mg protein, assuming 30% dry mass fraction, 50% protein fraction of dry mass, and a P/O ratio of 4.0 ATP per O₂, yields approximately 0.25 µmol ATP per mg protein per hour^48^. This value applies to metabolically active tissues; connective tissue and adipose tissue, which are metabolically less active, are excluded from this estimate.

---

**Figure S1:**
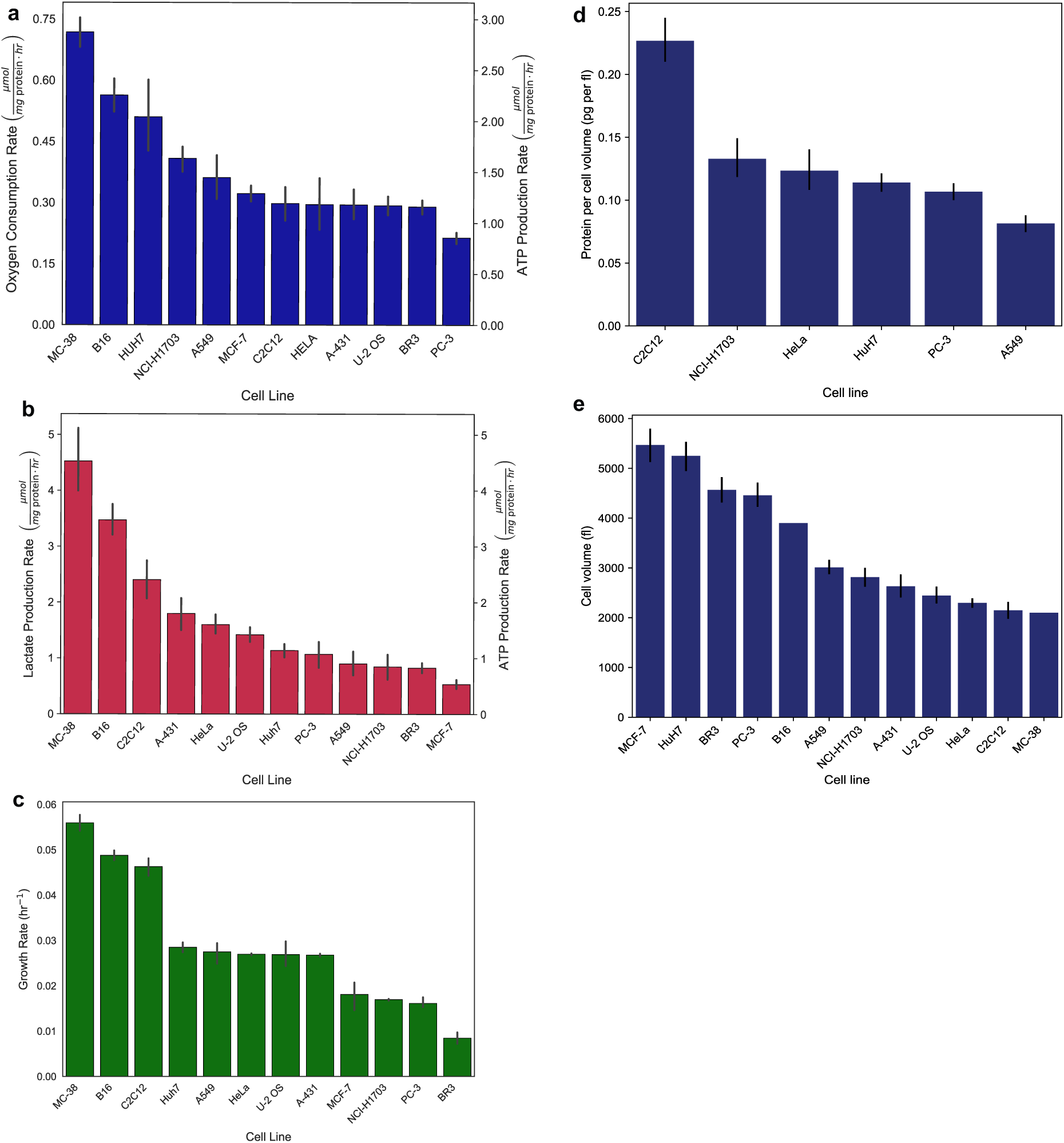
Oxygen consumption rate, lactate production rate, and growth rate of twelve mammalian cells lines. a, Oxygen consumption rates (µmol mg protein^-1^ hr^-1^), b, lactate production rates (µmol mg protein^-1^ hr^-1^) and c, growth rates (hr^-1^) of twelve mammalian cell lines. **d,** protein content per unit cell volume (pg per fl) for six mammalian cell lines. **e,** cell volume (fl) of twelve mammalian cell lines measured by Coulter counter. The error bars in represent the **95% confidence interval.**

**Figure S2:**
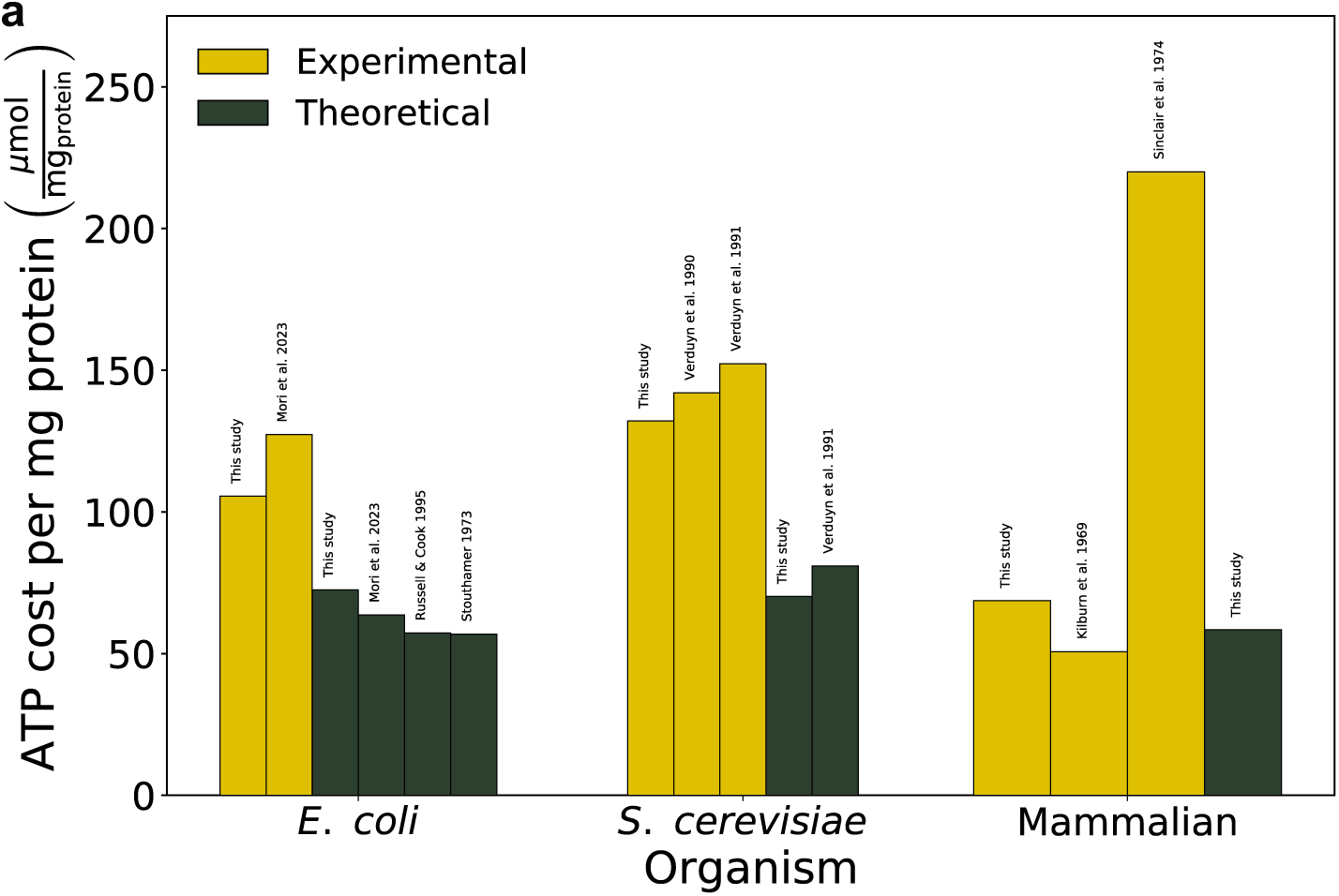
Comparison of ATP cost of biomass production across organisms. Experimental and theoretical estimates of ATP cost per unit biomass, expressed as µmol ATP per mg protein, are shown for *E. coli*, *S. cerevisiae*, and mammalian cells. Yellow bars denote experimentally measured ATP production or biosynthetic costs, whereas dark green bars denote theoretical estimates based on biomass synthesis. For each organism, values from this study are shown alongside representative literature estimates (Stouthamer 1973; Russell & Cook 1995; Kilburn et al. 1969; Sinclair et al. 1974; Verduyn et al. 1990, 1991; Mori et al. 2023). All values are normalized to cellular protein content to enable direct cross study comparison.

**Figure S3:**
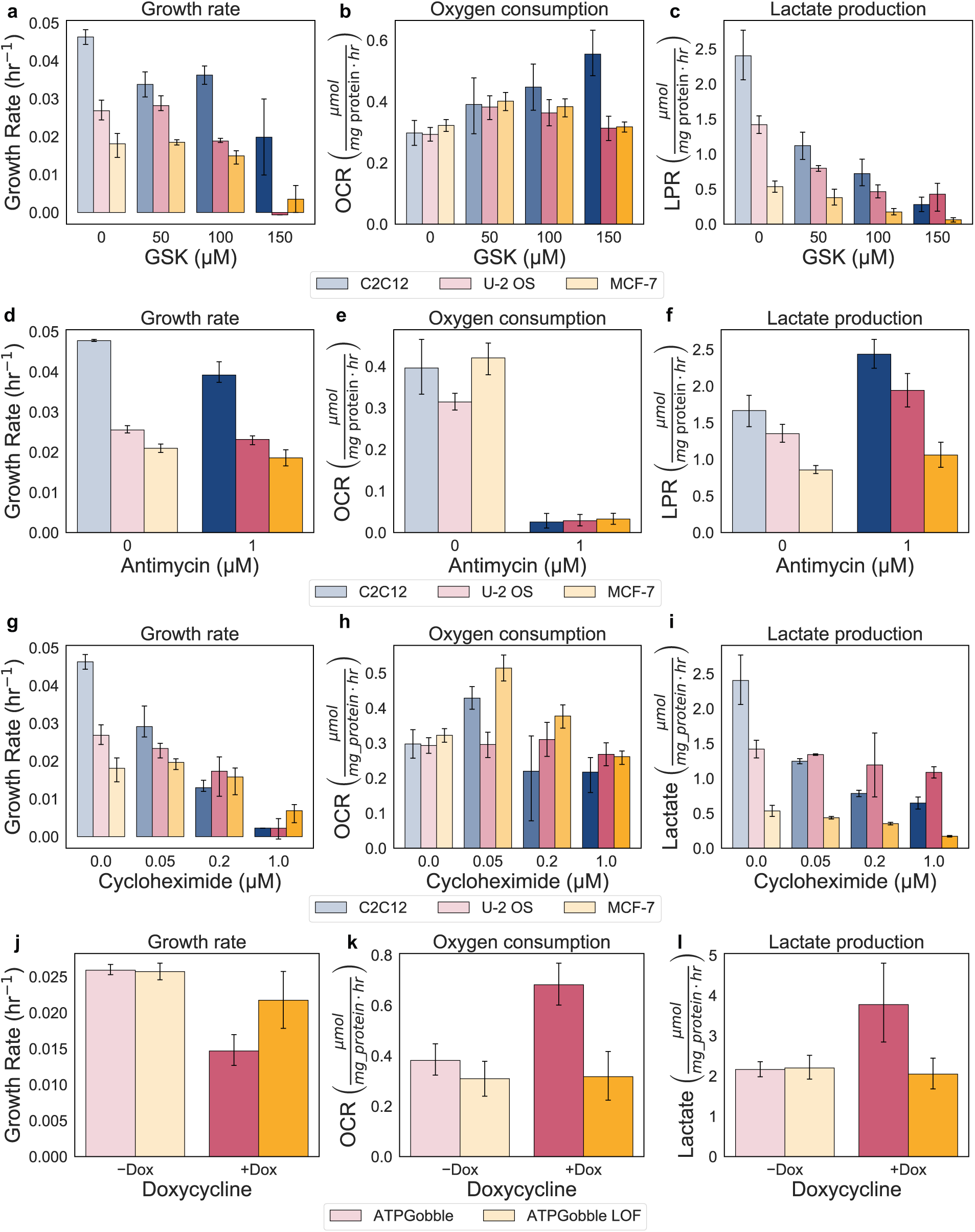
**Effects of glycolysis inhibition, respiration inhibition, biosynthesis inhibition, and ATPGobble expression on growth rate, oxygen consumption, and lactate production. a–c**, Dose response to the lactate dehydrogenase inhibitor GSK-2837808A (GSK, 0, 50, 100, 150 μM) in C2C12, U-2 OS, and MCF-7 cells, showing growth rate (hr⁻¹), oxygen consumption rate (OCR, μmol mg protein⁻¹ hr⁻¹), and lactate production rate (LPR, μmol mg protein⁻¹ hr⁻¹). **d–f**, Response to the Complex III inhibitor antimycin A (0, 1 μM) in C2C12, U-2 OS, and MCF-7 cells on the same three measurements. **g–i**, Dose response to the translation inhibitor cycloheximide (0, 0.05, 0.2, 1 μM) in C2C12, U-2 OS, and MCF-7 cells on the same three measurements. **j–l**, Response to doxycycline-induced expression of ATPGobble or its loss-of-function (LOF) variant in U-2 OS cells on the same three measurements. In all panels, error bars show 95% confidence intervals from 10,000 bootstrap resamples matched across measurements. Control reference points (untreated or −Dox) are shown as faded bars.

